# Correlated host movements can reshape spatio-temporal disease dynamics: modeling the contributions of space use to transmission risk using movement data

**DOI:** 10.1101/2024.04.16.589740

**Authors:** Juan S. Vargas Soto, Justin Kosiewska, Lisa I. Muller, Dan Grove, Dailee Metts, Mark Q. Wilber

**Affiliations:** School of Natural Resources, University of Tennessee, Knoxville, TN

**Keywords:** wildlife disease, host overlap, utilization distribution, animal movement

## Abstract

Despite decades of epidemiological theory making relatively simple assumptions about host movements, it is increasingly clear that non-random movements drastically affect disease transmission. To better predict transmission risk, theory needs to simultaneously account for how the environment affects host space use and how social dynamics affect correlation in space use. We develop new theory that decomposes the relative contributions of fine-scale space use and correlated movements to spatio-temporal transmission risk. Using analytical results, simulations, and empirical movement data, we show that even weak correlations can increase transmission risk by orders of magnitude compared to independent movement. Accounting for correlation is especially critical for pathogens with direct transmission or short environmental persistence. Our theory provides clear expectations for what has been observed empirically but largely ignored in disease models—movement correlation can reshape epidemiological landscapes, creating transmission hotspots whose magnitude and location are not necessarily predictable from spatial overlap alone.

## Introduction

Individual movement is a critical factor influencing wildlife disease dynamics (Dougherty et al., 2018; Manlove et al., 2022); Movement determines encounters with other individuals of the same species, other species, or pathogens in the environment (Martinez-Garcia et al., 2020; Das et al., 2023). These encounters are necessary for the transmission of infectious diseases, and efforts have sought to identify where they occur, how often, and how they are influenced by environmental and social drivers (Titcomb et al., 2021; Dougherty et al., 2022; Webber et al., 2023). Formally linking social factors, environmental factors, animal movement, contact, and pathogen transmission would improve our ability to predict and prevent outbreaks and represent a significant advancement for management of wildlife diseases. Nevertheless, understanding how these processes interact at an individual scale requires detailed movement information and theory to translate movement into an epidemiological context.

Most epidemiological theory is built upon the assumption of independent host movements, and there is little theory that quantifies how non-independent, correlated movements affect contact and transmission risk. Despite a large body of empirical work that quantifies how correlated and social movements can reshape contact and transmission landscapes (e.g., Kjær et al., 2008; Grear et al., 2010; Schauber et al., 2015), we lack models that isolate the role of social interactions on spatio-temporal force of infection (FOI, the risk of transmission experienced by a host per unit time). This limits our ability to ask questions like: how do non-independent movements affect spatio-temporal infection risk, compared to spatial overlap? Moreover, recent studies have shown that spatial transmission risk can be highly localized (Albery et al., 2021) and is not necessarily predicted by animal space use (Yang et al., 2023a). We hypothesize that non-independent animal movements can (at least partially) account for these observations and develop a modeling approach to rigorously test this hypothesis and systematically quantify the contribution of space use and correlated movements to spatio-temporal transmission risk.

Recent developments at the interface of movement and disease ecology leverage high-resolution animal tracking data to gain insight into contact among individuals and disease transmission (Richardson and Gorochowski, 2015; Wilber et al., 2022; Yang et al., 2023b). For example, movement-driven spatio-temporal infection risk (MoveSTIR) builds dynamic contact networks from movement data to estimate individual risk of infection across space and time (Wilber et al., 2022). MoveSTIR provides a theoretical foundation to translate contacts into the epidemiological currency of FOI. These studies have highlighted the importance of individual heterogeneity and temporal scale for epidemiology, particularly how indirect contact—individuals at the same place at different times—can significantly reshape contact and transmission networks (Richardson and Gorochowski, 2015; Yang et al., 2023b). Current approaches are nonetheless based on occurrence, rather than range, distributions (in the terminology of Alston et al., 2022) – meaning they only consider where animals were and not where they *potentially* could be. This approach makes it difficult to systematically link encounters with environmental drivers, and to predict how social or environmental changes affect contact and transmission.

Alternatively, spatio-temporal contact could be studied probabilistically, using utilization distributions (UDs). The UD represents the probability—transient or long-term (Tao et al., 2016)—of an organism using some area (Worton, 1989). The high spatial and temporal resolution of modern tracking data serves to build UDs based on biologically realistic movement models (Kranstauber et al., 2012; Fleming et al., 2014), and to link them with underlying resources (Potts and Börger, 2023). Additionally, combining individual UDs informs about pairwise interactions, by allowing to quantify home range overlap (Winner et al., 2018), or to estimate the expected location and rate of encounters (Noonan et al., 2021), which could serve to infer transmission risk (Godfrey et al., 2010; Godfrey, 2013; Noonan et al., 2021). Moreover, because UDs can be directly linked to environmental drivers of movement (Signer et al., 2017), they could be used for prospective analyses, to predict contact and transmission in novel environments, or to understand cascading effects of environmental and social perturbations from individual movement to population and landscape-level disease transmission.

Current contact metrics based on UDs focus only on direct interaction, ignoring temporal dynamics that are especially relevant for epidemiological processes. The Conditional Distribution of Encounters (CDE) (Noonan et al., 2021), for example, estimates local probabilities of encounter assuming that individuals move independently. While a useful simplification, social interactions like territoriality or gregariousness can invalidate this assumption (Manlove et al., 2018; Sah et al., 2018). In these cases, temporal correlations in space use could increase or decrease the probability of encounter expected given independent movement (Kjær et al., 2008; Schauber et al., 2015). Moreover, direct interactions do not necessarily equate to *epidemiological contacts*, which comprise contact formation, contact duration, pathogen acquisition, pathogen shedding, and pathogen decay. As some pathogens can persist in the environment for months or years (e.g. anthrax, chronic wasting disease–CWD), ignoring these processes could severely underestimate transmission risk (Wilber et al., 2022; Yang et al., 2023b; Richardson and Gorochowski, 2015).

Here, we develop a model we refer to as Probabilistic MoveSTIR (PMoveSTIR) to estimate epidemiological contact and expected FOI across space and time from UDs and correlated animal movements. We derive a general model that links transient, spatially heterogeneous UDs to transmission, and provide modifications to consider stationarity in space or time. Deriving analytical results and applying PMoveSTIR to simulated data, we demonstrate the sometimes sizable importance of correlated host movements on transmission risk, indicating that ignoring social drivers of contact could severely bias epidemiological inference. Using data on movement of white-tailed deer (*Odocoileus virginianus*), we show that correlated movements can increase potential FOI by orders of magnitude for a hypothetical directly transmitted pathogen but are relatively unimportant for a hypothetical pathogen with long persistence times. Moreover, non-independent movements can yield highly heterogeneous transmission landscapes, supporting the emerging idea that finescale patterns of transmission dynamics are common in wildlife (Albery et al., 2021) and are contributed to by correlated host movements. By precisely quantifiying the two fundamental components of non-random movements, fine-scale space use and correlation in movement, PMoveSTIR is a key next step for developing predictive models that link movement data to spatio-temporal infection dynamics on real landscapes.

## Methods

### Model development - Linking utilization distributions to transmission through PMoveSTIR

PMoveSTIR builds on the MoveSTIR model (Wilber et al., 2022) and formally links UDs, direct and indirect contacts, correlated movements, and spatial estimates of FOI. Essentially, we want to know, for two individuals *i* and *j* moving and interacting across a landscape, what the FOI *i* experiences from *j*, across space and time is.

As in MoveSTIR, we assume that transmission happens by an infected host depositing pathogen into the environment and another host picking that pathogen up. Deposition and acquisition can represent a range of processes, from coughing and inhaling in a matter of seconds, to depositing parasite eggs or larvae in the environment and another individual consuming these days or weeks later. Considering transmission through deposition and acquisition clearly links direct and indirect transmission along a continuum (Wilber et al., 2022), and it encompasses standard density-dependent transmission as a special case (Cortez and Duffy, 2021).

In PMoveSTIR, “contacts” occur if both individuals visit a given location *x*, which could be a habitat patch or a grid cell. Locations *x* do not overlap so the sum of their areas equals some total area over which individuals can move (see Appendix 1 for a derivation when *x* is a point, not an area). Furthermore, we assume that the likelihood of contact is uniform within *x*, consistent with a so-called top-hat encounter function (Gurarie and Ovaskainen, 2013; Wilber et al., 2022).

Given these assumptions, we define the pairwise FOI host *j* exerts on host *i* in location *x* at time *t* as (Wilber et al., 2022)

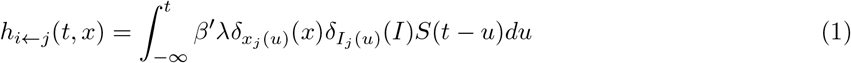

where *λ* is the pathogen deposition rate, 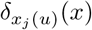is an indicator variable that is one if host *j* is in location *x* at time *u* and zero otherwise, 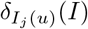 is an indicator function that is one if host *j* is infected at time *u* and zero otherwise, and *S*(*t* − *u*) is the probability that pathogens deposited at time *u < t* are still transmissible at time *t* (see Wilber et al., 2022, for a full derivation). The term *β*^*′*^ is the rate at which host *i* acquires pathogen within location *x* and can be re-written as 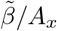, where 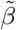 can be considered a “search efficiency” term, with units area/time (e.g., *m*^2^*/day*), and *A*_*x*_ gives the area of location *x* (e.g., 100 *m*^2^). Therefore, the total acquisition rate scales with the area in which contact can occur; in larger areas, the acquiring host would have to search for longer to encounter an infectious dose, reducing the total acquisition rate and the corresponding FOI. Moving forward, we assume that the depositing host is always infected, and shedding pathogen infectious stages at a constant rate. This assumption is equivalent to building a contact network and also represents the structural form of FOI needed to compute pathogen invasion thresholds (Wilber et al., 2022).

Considering probabilistic space use (i.e., we know where an individual is at a given time and location with some probability), we can re-write equation 1 as

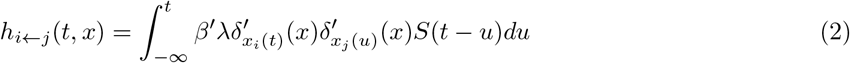

where 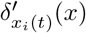 is a random variable that specifies whether host *i* is in location *x* at time *t* (defined equivalently for *j*). This means that *h*_*i*←*j*_(*t, x*) is also a random variable, and we can express its expected value as

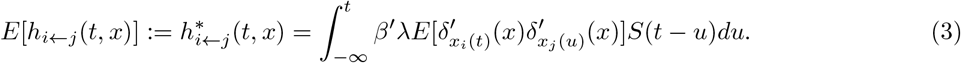

Interpreting this expectation, we are asking: if we simulated some movement process thousands of times, what is the probability that host *i* is in location *x* at time *t*, and host *j* was in *x* at a previous time *u*?

### Linking equation 3 to utilization distributions

For two random variables *Y* and *Z, E*[*Y Z*] = *E*[*Y*]*E*[*Z*] + *Cov*(*Y, Z*). We can therefore write equation 3 as

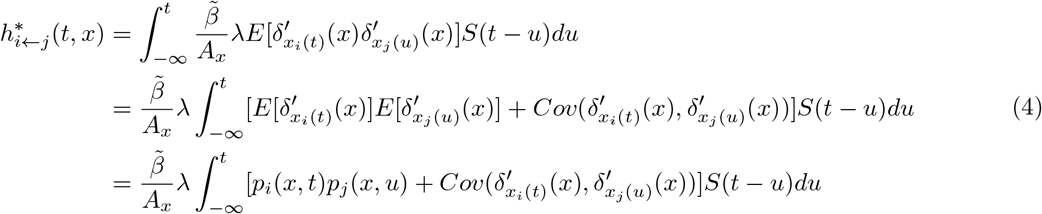

where we use the fact that the expectation of an indicator variable is a probability (Grimmett and Stirzaker, 2001). The terms *p*_*i*_(*x, t*) and *p*_*j*_(*x, u*) give the probabilities that host *i* and *j* are in location *x* at times *t* and *u*, respectively, and can also be written as 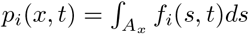 where *f*_*i*_(*s, t*) is the probability density of host *i* using the point *s* at time *t* and the integral is over the area *A*_*x*_ (defined equivalently for host *j*). Thus, we have obtained an equation that links the transient utilization distributions *f*_*i*_(*s, t*) and *f*_*j*_(*s, u*) with the spatio-temporal FOI. This general PMoveSTIR formulation accounts for heterogeneity in space and time, but the model can consider other scenarios, such as uniform space use, or UDs stationary in time (Fig. 1, Appendix 2).

**Figure 1:**
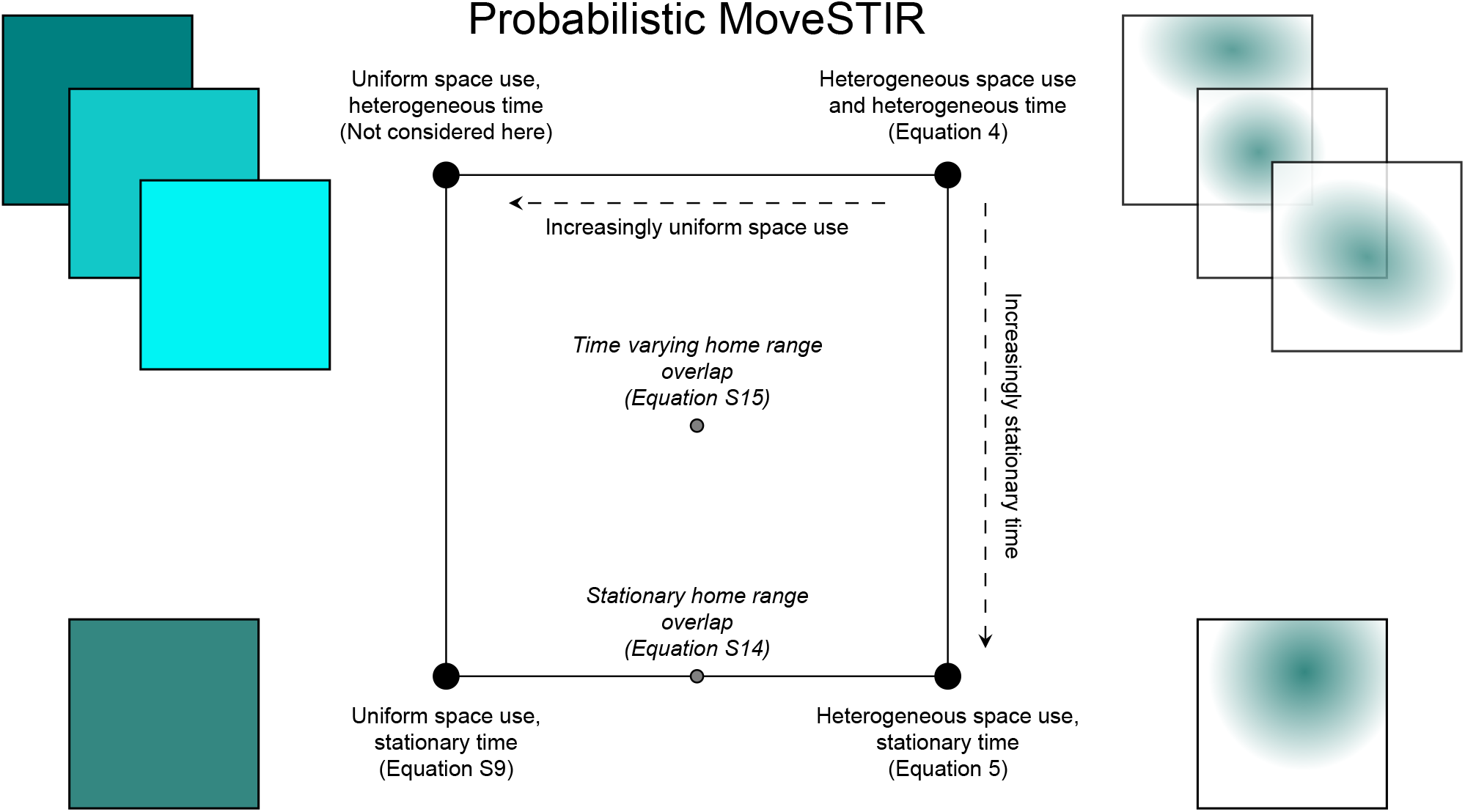
Conceptual figure describing the model developed in this manuscript: probabilistic movement-driven modeling of spatio-temporal infection risk. PMoveSTIR can be thought of as square where the dimensions represent heterogeneity in space and time. The upper right-hand corner is the most general case: heterogeneous space use by hosts, and movement dynamics that are not statistically stationary. As space becomes increasingly uniform or movement becomes more statistically stationary, PMoveSTIR reduces to the upper-left hand corner or the lower-right hand corner, respectively. We primarily focus on the lower-right hand corner in this manuscript. When space use is uniform and movement is statistically stationary, we are in the lower-left hand corner and recover mass action transmission as a special case (Appendix 2; assuming host movements are uncorrelated).

For example, when UDs are stationary in time, and pathogen decays exponentially in the environment at rate *ν*, equation 4 becomes (derivation in Appendix 2)

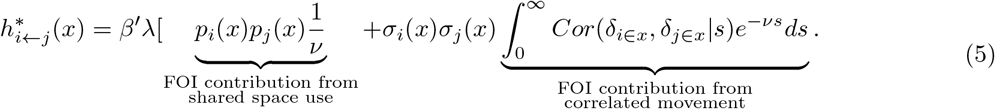

where 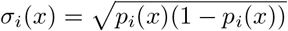 is the standard deviation in probability of host *i* using location *x*. Importantly, this shows that the expected FOI between two individuals is a result of 1) shared space use, and 2) correlation in host movements across different time lags.

### Analytical and simulation insight into correlated movement and FOI

Leveraging PMoveSTIR, we used analysis and simulation to ask: how much can correlated movements affect spatio-temporal infection risk for directly and indirectly transmitted pathogens relative to spatial overlap? First, we used PMoveSTIR to derive equations that explicitly quantify how much correlation can augment or reduce force of infection due to direct or indirect contact compared to random movement. Second, we used simulations to explore how temporally correlated movements affect transmission, and how the pairwise FOI estimate depends on epidemiological factors such as contact distance and pathogen decay. Additionally, empirically estimating the correlation component in equation 5 depends on the amount of data available. Therefore, we also explored the implications of tracking duration (the total time an animal is tracked) on the estimated FOI. For the simulations, we focused on the lower-right corner of Fig. 1 (equation 5), where we assumed statistical stationarity in movement. The process for calculating the FOI across the landscape is summarized in Fig. 2.

**Figure 2:**
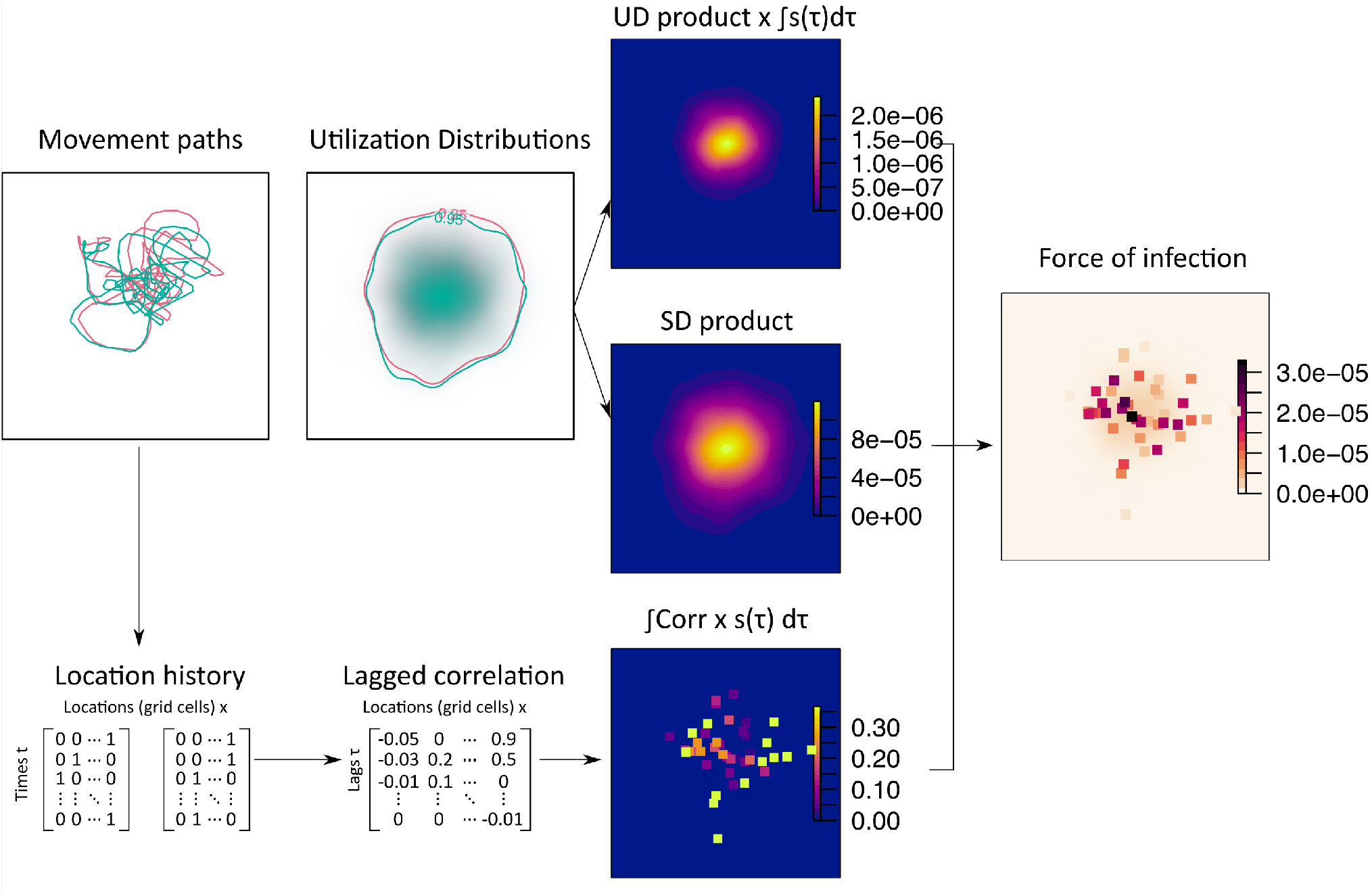
Flow diagram for calculating spatial force of infection (FOI) from movement data using PMove-STIR. Starting from position data, we estimate individual utilization distributions, and use them to estimate the product of the UDs and their standard deviations (SDs). In addition, individual position histories are used to estimate the pairwise temporal cross-correlation in space use at each cell (e.g., the correlation term in equation 5). All three elements are combined and scaled by epidemiological parameters to obtain a spatially explicit, pairwise, directional FOI. Note that here and in the manuscript we do not spatially interpolate the correlation surface. We therefore provide a conservative view of how correlation contributes to landscape level FOI.

We simulated 20 replicate movement tracks for each combination of parameters. In every simulation, two individuals moved around established home ranges according to an Ornstein-Uhlenbeck process; i.e., they move randomly but are attracted back towards the center of their home range as they drift away. To create different levels of correlation, we modified the initial simulated tracks using a convolution approach with a social interaction kernel (Scharf et al., 2018). This method accounts for constant or temporally varying attraction between pairs of individuals. For our purposes we assumed attraction strength was constant in time, but varied across pairs from 0 (completely independent movement) to 1 (joint movement). For every scenario, we fitted continuous-time movement models (CTMM) to the simulated tracks, and estimated individual UDs using autocorrelated kernel density estimation (AKDE, Calabrese et al., 2016). We estimated the UDs on a grid of square cells, where the cell side *d* is the approximate threshold contact distance for epidemiological contact.

We used the UDs to calculate the product of the probabilities of use (*p*_*i*_(*x*)*p*_*j*_(*x*)) and the product of their standard deviations 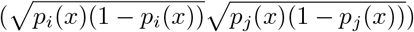 for each grid cell. Both products are symmetrical for every pair of individuals. The lagged correlation term in equation 5 is calculated based on the position history for each individual at locations that both visited (locations that only one or neither individual visited have a correlation of zero). The position history is a binary vector that specifies whether each individual was present (1) or absent (0) at location *x* at time *t*. The order of visits matters, so these correlation values can be asymmetric between individuals. Correlations can appear spuriously even in the absence of a true interaction, particularly for time series that are autocorrelated. To address this issue, we performed a pre-whitening step to remove the potential effects of autocorrelation on the estimated cross-correlation.

Pre-whitening consists of fitting an autoregression model to one of the series, and filtering both time series using the coefficients from that model (Dean and Dunsmuir, 2016). Additionally, we retained only correlation values deemed significantly different from 0, i.e. correlations with absolute values greater than a threshold of 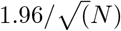, where *N* is the length of each time series (Dean and Dunsmuir, 2016). All other values were set to 0 as they are considered random noise. We then scale each correlation value by the exponential pathogen survival function *S*(*s*) = *e*^−*νs*^, where *s* is the lag corresponding to each cross-correlation (between 0 and *t* − *dt*), *dt* is the (constant) time lag between observations, and *ν* is the decay rate of the pathogen in the environment.

Substituting the terms in equation 5 and scaling by the epidemiological parameters 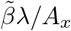 we obtain the per-cell FOI. Through these simulations we explore how the expected FOI is influenced by correlation in space use, home range overlap, pathogen decay rate, and contact distance.

### Empirical application - White-tailed deer

To test the role of space-use and correlated movements on potential transmission risk in a real system, we applied the PMoveSTIR model to GPS-tracking data for five white-tailed deer from Ames Plantation, Tennessee, USA (one adult male and four adult females). Deer were captured and equipped with GPS collars that recorded fixes every 30 minutes (Lotek LiteTrack Iridium 420, Newmarket, Ontario, Canada; IACUC # 2850-1021 from the University of Tennessee). All individuals used in this study were captured at the end of March 2023 but we only included movement data from May to June in this study, to eliminate any effects of capture. As for the simulations, we used CTMM and AKDE to estimate individual UDs (Calabrese et al., 2016). We used the CTMMs to interpolate the positions to regular 10-minute intervals, to fill in missed positions, and to get the same timestamps across individuals (Yang et al., 2023b). Assuming a threshold contact distance of *d* = 10m, we divided the landscape in a grid of 10×10m cells.

We modeled two hypothetical pathogens. The first pathogen had a relatively short persistence time in the environment, surviving for an average of 1 hour (*ν* = 1 h^−1^). In this case, transmission is largely direct and this might represent a pathogen like SARS-COV-2, which can infect and transmit between white-tailed deer (Hale et al., 2022). The second hypothetical pathogen had a long persistence time, remaining viable for over a year on average (*ν* = 0.9 yr ^−1^). In this case, transmission is largely indirect and might reflect a pathogen like CWD, which can transmit directly and indirectly between deer and can persist for years in the environment (Saunders et al., 2012). The *β* and *λ* parameters are scalars in PMoveSTIR and do not affect any relative comparisons, so we set them both to unity. As in the simulations, we pre-whitened and filtered correlation values to remove potentially spurious correlations. We used these data to explore how differences in overlap across home ranges and correlated movement influence the expected FOI with real animal trajectories.

## Results

### The importance of correlated movements on FOI – analytical results

Consider the situation where movement is statistically stationary and hosts use space uniformly (equation S9). For illustrative purposes, consider two hosts moving together across some area *A*_*tot*_. We assume that 1) hosts spend *η* time units within a habitat patch/grid cell before moving to the next one, 2) the pathogen survival function *S*(*s*) is a step function with a survival probability of one when lag *s* ≤ *πη* and zero when *s > πη*. The term *πη* gives the time the pathogen survives in the environment as a function of host residence time, where *π* ranges from near zero for directly transmitted pathogens to some arbitrarily large number for pathogens with long environmental persistence. In this scenario, transmission can only occur while both animals are in the same patch. With these assumptions, we can rewrite equation equation S9 as

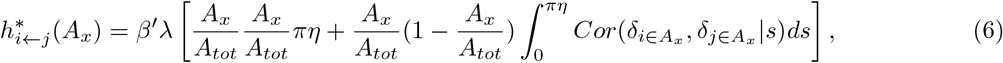

recognizing that 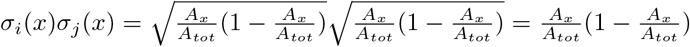 when both hosts are using space uniformly.

For hosts that are moving together, 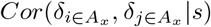 will be exactly unity when lag *s* = 0 and near unity when lag *s* is near zero. When pathogens are strictly directly transmitted, *π* is also small and if *πη << η* then we can reasonably approximate 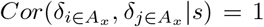 for s from 0 to *πη*. We can then write equation 6 as

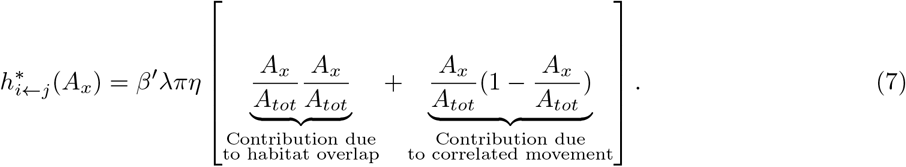

The relative contribution of correlation in movement with respect to the contribution due to habitat overlap is simply (1 − (*A*_*x*_*/A*_*tot*_))*/*(*A*_*x*_*/A*_*tot*_) = *A*_*tot*_*/A*_*x*_ − 1. Thus, PMoveSTIR allows us to put intuitive bounds on the importance of correlated movements for direct transmission risk. As the area *A*_*x*_ in which an epidemiological contact can occur gets smaller relative to the total area in which the hosts are moving *A*_*tot*_, correlated movement can have an orders of magnitude larger contribution to direct transmission FOI than habitat overlap (Fig. 3). This finding makes intuitive sense but is almost never considered in epidemiological models. If the area of potential contact is small and hosts are moving randomly, there is a very low chance that hosts will be there together at the same time. Correlated movements significantly increase this chance, even when correlation is low (e.g. 0.1). The contribution of correlation to FOI can be 10 times greater than spatial overlap if the area of contact is small (e.g., *<* 5% of the total area; Fig. 3). Large correlations across multiple areas result in a significantly greater FOI across the entire landscape.

**Figure 3:**
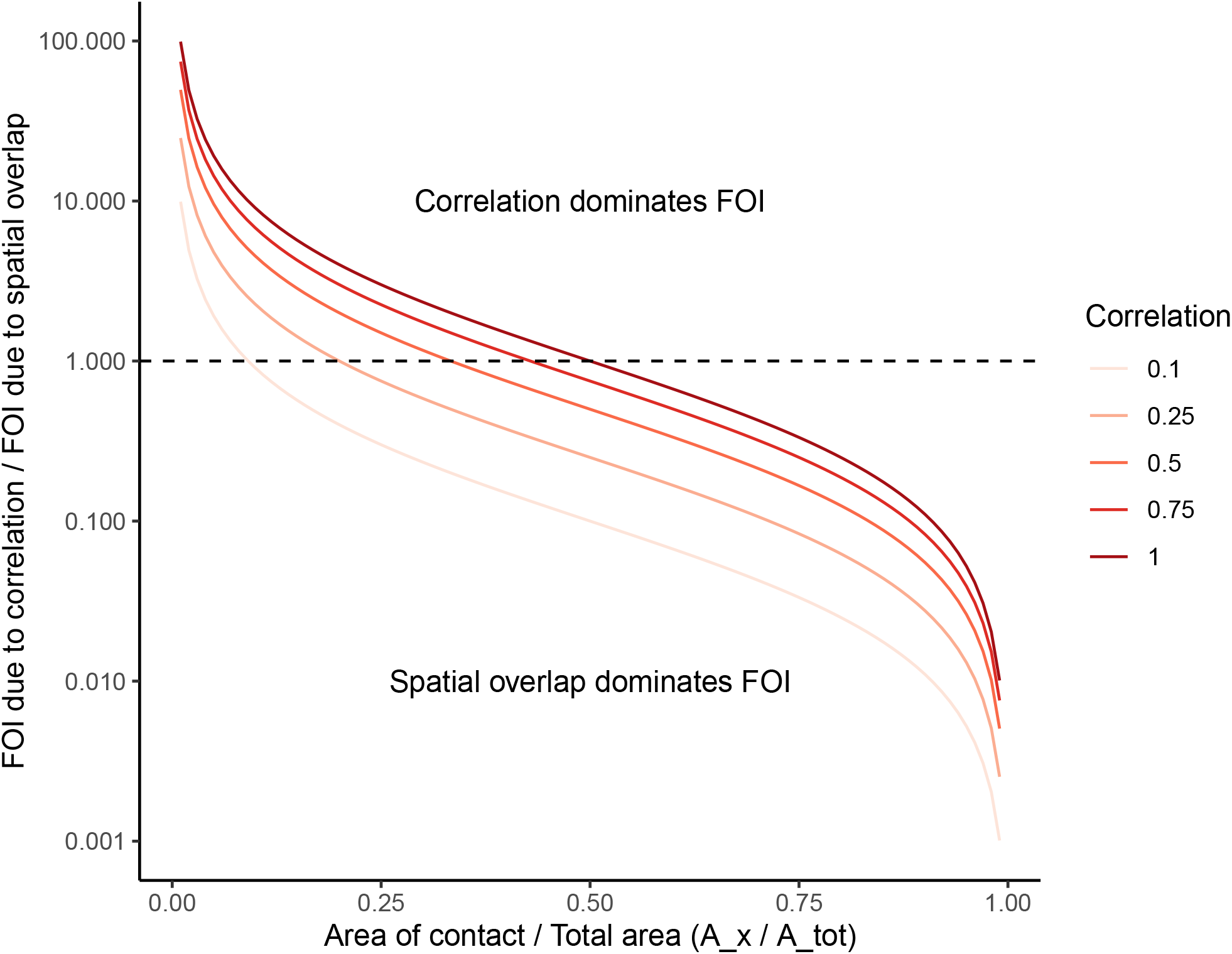
The relationship between the relative contribution of correlation in movement and spatial to pairwise force of infection (FOI) and the relative area of contact. This analytical relationship is derived in equation 7 assuming direct contact. When the relative area of contact is small relative to the total area over which animals can move (e.g., *<* 10%), even weakly correlated movements can significantly increase FOI relative to spatial overlap.

In contrast, when *A*_*x*_ approaches *A*_*tot*_ (or more generally when the probability of using a particular area is very high), the importance of correlated movement relative to habitat overlap becomes minimal (Fig. 3). For example, if two hosts use a particular area together because of high resource availability, then it does not matter for FOI if social factors are leading to additional correlated movement. This result extends beyond epidemiological contexts and shows when correlated movements can significantly alter contact risk based on metrics such as home range overlap or CDE. In Appendix S3, we expand this example to examine the effects of correlated movement on FOI given indirect transmission.

## Simulation study

### Effect of correlated movement on pairwise FOI

In our simulations, greater interaction strengths led to higher overlap among home ranges, but there were also pairs moving independently with similarly high (*>*0.9) overlap. These scenarios allowed us to compare the FOI while teasing apart the effects of spatial overlap and temporal correlation in space use.

The FOI was higher for pairs with higher interaction values (i.e. with stronger attraction), in part due to the resulting increase in spatial overlap. However, we saw an even greater increase in FOI when we accounted for temporal correlation, and the difference between the two increased with higher interaction strengths (Fig. 4a). For very similar trajectories (interaction*>*0.9), the FOI with correlation was more than ten times the FOI estimated without correlation, and there was a more than 100-fold difference for perfectly overlapping trajectories.

**Figure 4:**
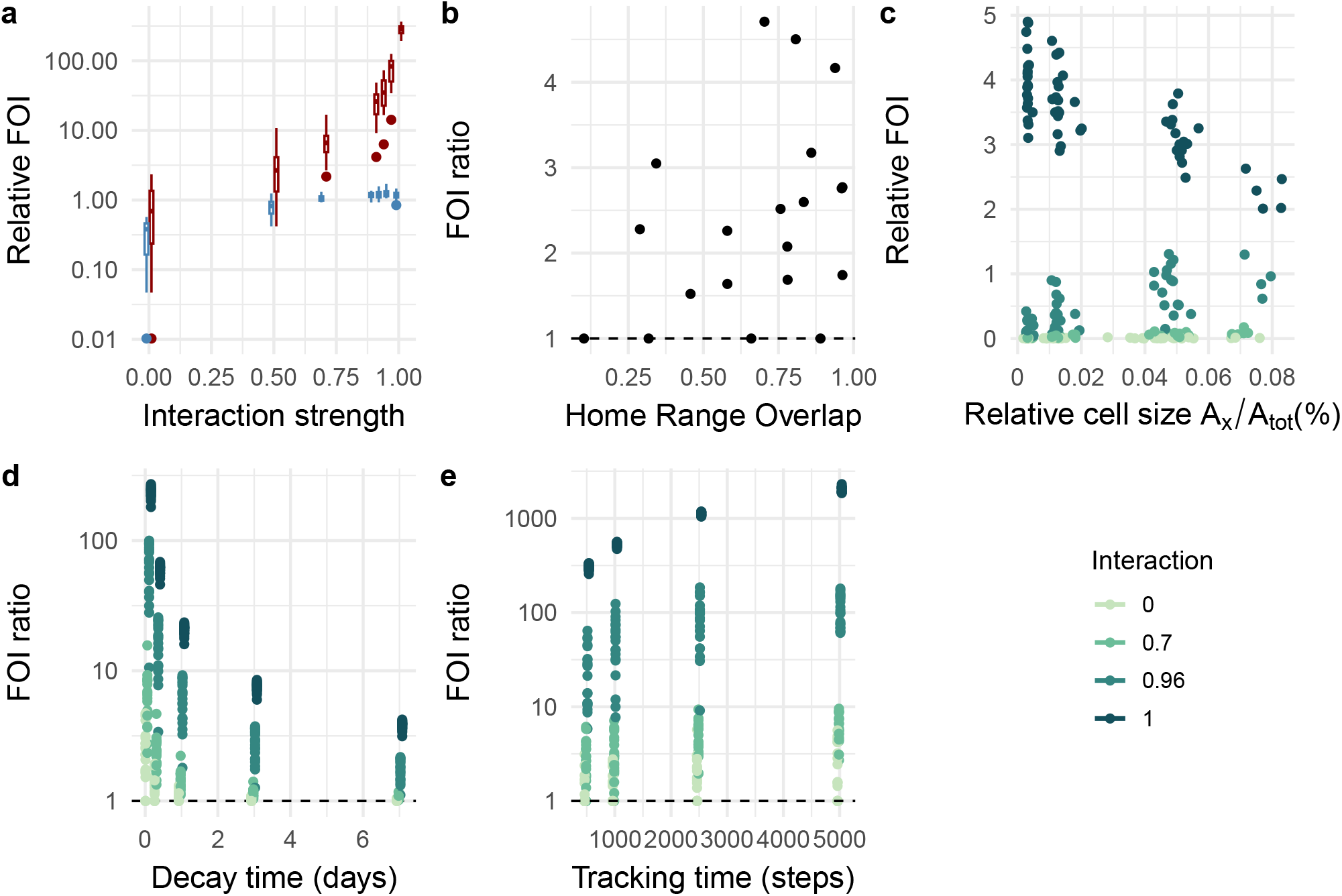
Analysis of simulated movement data show how the force of infection (FOI) varies depending on strength of attraction between individuals and epidemiological parameters such as the pathogen decay rate, the threshold distance that defines a contact, and the data availability. **a)** FOI generally increases as a function of attraction strength, but the estimated values are greater when correlation is considered (red boxplots) than when only spatial overlap is considered (blue); **b)** The estimated FOI as a function of home range overlap (as measured by the Bhattacharyya coefficient) shows how correlation can have an apparent effect even for animals that move independently; **c)** Longer epidemiological contact distances can increase, or have no effect on the estimated FOI, depending on the interaction strength; **d)** The contribution of correlation to the overall FOI decreased with longer decay times; **e)** The FOI ratio increases with longer time series as correlation information becomes available for a larger proportion of the space used. A FOI ratio greater than one indicates that correlated movement is increasing FOI relative to spatial overlap and a ratio less than one indicates that correlated movement is decreasing the FOI. In c-e, darker shades indicate stronger attraction between individuals.

The FOI was generally greater when we accounted for correlation; in only 19% of cases the estimates were equal with or without correlation, and in no instance was there a decrease. This effect is to be expected as we only included attraction in the simulations. We found that correlation could increase the estimated FOI even for pairs that moved independently. While this was generally rare (86% of cases had less than 20% increase), there were instances where the estimated FOI was more than three times greater. Larger effects were more likely for pairs with a high degree of home range overlap (Fig. 4d).

While the expected correlation is 0 when individuals move independently, joint space use could lead to apparent correlations, especially when there are few observations in a given cell. These correlations can persist after pre-whitening and removing the majority of statistically spurious correlations. Apparent correlations are important to consider when analyzing empirical data where the true correlation and the degree of attraction/avoidance among individuals are unknown. Temporal reshuffling methods such as those proposed by Spiegel et al. (2016) can be a useful tool to establish the baseline “apparent” correlations that appear for non-socially interacting individuals.

### Influence of epidemiological parameters

The FOI was inversely proportional to decay rate, meaning transmission risk is greater for pathogens that can survive longer in the environment, all else being equal (Fig. 4b). However, the increase in FOI due to correlated movement is greatest for pathogens with short survival times. While longer pathogen survival times increase the probability of indirect epidemiological contact, the expected rate of encounter at longer lags approximates the expected rate of encounter under independent movement, so correlations at long lags have only a small modifying effect on the expected FOI (Fig. 4e).

The effect of contact distance on FOI depended on interaction strength and the effect of correlation (Fig. 4c). Consistent with our analytical results, when animals are highly correlated in their movements (i.e., have a high interaction strength) the importance of correlation is diminished as contact distances increase. However, our simulations also show that for non-perfectly overlapping individuals, longer contact distances can increase the probability of finding two individuals in the same cell sufficiently to offset the inherent reduction in transmission in larger areas (remember, 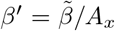), therefore increasing the relative contribution of correlation to FOI for trajectories that are already close in space (Fig. 4c).

### Influence of data availability

With longer time series, the number of cells that both animals visited (where correlation is calculated) increased linearly. This resulted in greater overall FOIs (although local correlation values could decrease) with more tracking data (Fig. 4e).

Since we did not include any environmental drivers of movement, the expected correlation is uniform across the landscape. The cumulative correlation, however, is not related to the product of either the UDs or the standard deviations. This result highlights the challenge of predicting an expected correlation based on currently used metrics of movement and space use.

### Empirical example: white-tailed deer

The total potential FOI varied more than six orders of magnitude across pairs of deer, partially due to differences in spatial overlap, which ranged between 0 and 90% (Bhattacharyya coefficient; Fig. 5). While higher overlap generally correlated with higher FOIs, we estimated significantly higher values when we accounted for correlation in movement, and the difference was not necessarily related with overlap (Fig. 5a-c). For the pair with the highest spatial overlap, the FOI was more than ten times greater when we accounted for correlation. Temporally reshuffling the trajectories of these individuals following Spiegel et al. (2016) confirmed that this increase in FOI due to correlation was far beyond what we would expect from any spurious, “apparent” correlations that could arise in independently moving hosts (as identified in our simulations). For all pairs considered, ignoring correlation could result in between 5% and 96% underestimation of FOI. These effects of correlation on FOI were strongest for pathogens with a fast decay rate. For a pathogen like CWD that can persist for very long in the environment, the FOI was virtually identical with or without correlation in host movement (Fig. 5c).

**Figure 5:**
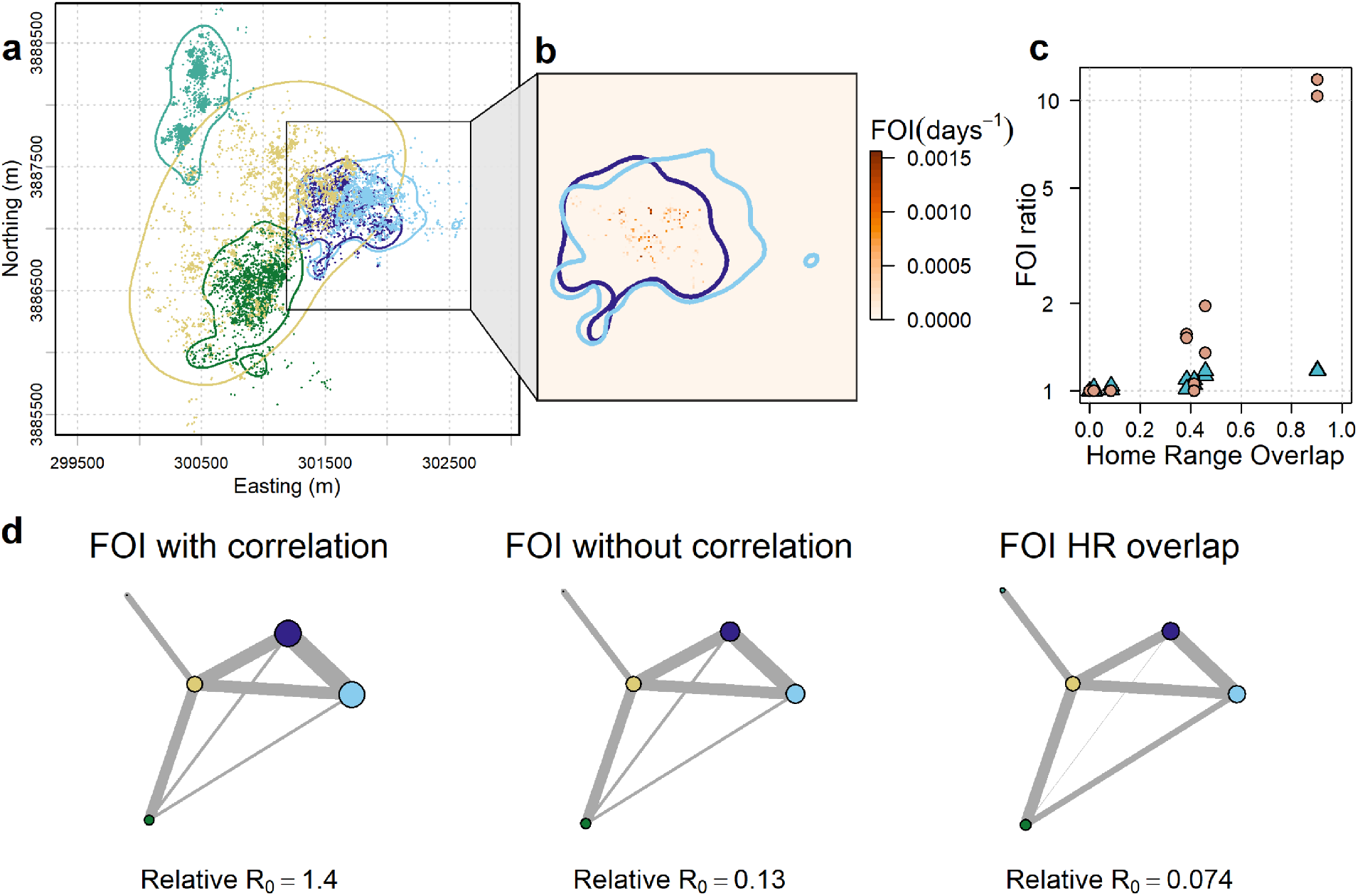
Application of the PMoveSTIR framework to movements of white-tailed deer in TN, USA. **a)** Home ranges of five individuals with different degrees of overlap. The points show the GPS locations, and the lines are the boundaries of the home ranges, estimated as the region that contains 95% of the utilization distribution density. **b)** Detail of the home ranges of the two individuals that overlapped the most. The color surface shows the estimated FOI, highlighting how accounting for temporal correlation creates a heterogeneous surface of disease transmission risk with distinct hotspots. **c)** The ratio of FOI values calculated with versus without correlation shows increased relevance of correlation with higher home range overlap. This effect is greater for pathogens with short environmental persistence like SCV2 (orange dots) than for pathogens with long persistence like CWD (blue triangles). **d)** Transmission networks created with the PMoveSTIR framework including correlation (left), without correlation (middle), and using the home range overlap (right). The size of the nodes represents the cumulative FOI experienced by each individual, and the width of the edges represents the pairwise FOI. We show only one direction here, but values were similar in both directions within a pair. Sizes and widths have the same scale in all three networks. *R*_0_ values should only be interpreted with respect to each other and not in absolute value.

Given the localized nature of interactions, the changes in overall FOI are caused by increases at particular cells. Consequently, the estimated FOI is heterogeneous across the landscape (Fig.5b). These effects at the local and individual scale affect population-level pathogen transmission. For example, given a simple Susceptible-Infected-Recovered model and short pathogen persistence time in the environment, the *R*_0_ on our network of five individuals was more than ten times greater when we accounted for correlation (Fig. 5d). Thus, we would predict more explosive transmission through a network that considers fine-scale space use and correlation compared to one that assumes independent movement.

## Discussion

We develop a new model, PMoveSTIR, that disentangles the contributions of spatial overlap and correlated movements to spatio-temporal contact and transmission risk. Our results provide clear expectations for what has been previously observed empirically but largely ignored in movement and disease models – correlation in movement can reshape epidemiological landscapes, leading to hotspots of transmission whose magnitude and location are not necessarily predictable from models of joint space use, in particular for directly transmitted or short-lived pathogens.

PMoveSTIR provides a clear, quantitative guide to assess when fine-scale, temporally synchronous movement data are necessary for capturing disease dynamics and when coarser scale, asynchronous data focused strictly on UD estimation are sufficient. In particular, we show that correlation in movement and synchronicity can be critical aspects of transmission for faster-paced pathogens, i.e. pathogens with short environmental persistence and transmission driven by short-term contacts (cf. Dougherty et al., 2018; Manlove et al., 2022), such as canine distemper virus, rabies, or SARS-COV-2 (SCV2). SCV2 is of particular public health interest and recent studies have shown that it can infect a wide-range of wildlife hosts, including white-tailed deer (Palmer et al., 2021; Hale et al., 2022). There is also evidence that white-tailed deer can transmit SCV2 between each other in controlled experiments and in the wild (Martins et al., 2022; Hale et al., 2022), suggesting SCV2 can successfully invade, spread, and persist in wildlife. In our analysis of empirical white-tailed deer movements, we found that ignoring correlation in movements and focusing only on patterns of space use led to a ten-fold change in *R*_0_ for a SCV2-like pathogen on even a simple network, with significant implications for pathogen spread. In contrast, our simulation and empirical results show that correlation may only have marginal effects on transmission of pathogens with longer persistence times like CWD and anthrax, and joint space use would be largely sufficient for understanding local transmission risk.

Correlations in space use that are not accounted for by UDs typically reflect social interactions, for example herding animals, parents with their offspring, or mates temporally moving together (Scharf et al., 2016; Yang et al., 2021). White-tailed deer female groups, for example, have high social affinity. Thus, pairs of deer with equivalent habitat overlap have substantially higher contact rates when both individuals are within the same social group (Schauber et al., 2007; Kjær et al., 2008; Schauber et al., 2015; Grear et al., 2010). This is likely the case in our empirical study; the pair with the greatest home range overlap and potential FOI was a mother–daughter pair. The overlap and interaction among them is consistent with the rose petal hypothesis (Porter et al., 1991), in which the home ranges of offspring radiate around the home range of their parent. The potential FOI in this case was over 10 times greater than for other pairs, despite similar degrees of habitat overlap. Importantly, 91% of the total FOI (for a hypothetical directly transmitted pathogen like SCV2) would be missed if we only considered overlapping UDs for these individuals (e.g., using a metric like CDE Noonan et al., 2021). While the importance of social interactions for contact rates has been documented previously in white-tailed deer (Grear et al., 2010; Schauber et al., 2015), PMoveSTIR allows us to directly quantify the contribution of these social interactions to the FOI and move from qualitative conclusions to movement-informed quantitative models of disease invasion.

In practice, however, correlations influencing FOI may arise from fine-scale temporal patterns of resource use unrelated to social interactions. For instance, consider two individuals independently visiting a common resource at similar times (e.g., a watering hole, VanderWaal et al., 2017). If the resource is highly localized in space and time, the joint, stationary UD would not necessarily predict this as an area of high contact and transmission risk. Rather, observed contacts in this area would be associated with correlation in movement and not long-term, predicted space use. Alternatively, temporally varying UDs (i.e., the top right corner in Fig. 1) would correctly capture that the joint occurrences were a result of spatio-temporal patterns of space use, reducing the relative importance of correlations on FOI. Unfortunately, non-parametrically estimating transient UDs at fine temporal scales with limited data points is not often feasible. Therefore, while the correlation term of PMoveSTIR will remain crucial to capture epidemiologically important spatial correlations in movement, it does not always provide an exact separation between spatial and social factors driving transmission risk.

By linking movement data and inferred contact with underlying environmental factors, a broad goal of PMoveSTIR is to create an epidemiological risk landscape independent of geographical location (Merkle et al., 2018; Manlove et al., 2022). This landscape could enable out-of-sample prediction of transmission risk, for example to forecast future disease dynamics in the study population, project population-wide disease spread as information on more individuals becomes available, or quantify transmission risk for other populations in similar environments. Our model provides a generalizable theoretical foundation to perform similar analyses across different host-pathogen systems, and it can be integrated with any UD estimation method (Signer et al., 2017; Merkle et al., 2018; Michelot et al., 2020; Potts and Börger, 2023). In addition to movement covariates, PMoveSTIR can also incorporate spatially and temporally heterogeneous epidemiological parameters, for example pathogen survival rates that vary between habitats and seasons (Daversa et al., 2017), or spatially localized shedding (Weinstein et al., 2018).

A key challenge for making out-of-sample predictions is empirically estimating the correlation surface. While this term is well-defined in theory (equation 5), finite movement data can inherently only provide a patchy view of the true correlation surface (e.g., Fig. 2). Our simulations showed that when animals were experiencing known social attraction at a time lag of zero, empirically estimated spatial correlations underestimated the contribution of correlation to FOI. This underestimation is not surprising because, without interpolation, we are limited to analyzing correlations where joint occurrences were observed and not where they potentially could have occurred. We did not use spatial interpolation tools (e.g., Kriging from a spatially-explicit statistical model, Clough-Tocher 2D interpolators, etc.) so our estimates provide a conservative view of how correlation could reshape epidemiological landscapes. To adequately predict transmission landscapes out-of-sample, future work will need to develop robust approaches for estimating and predicting the entire correlation surface.

Predicting epidemiological landscapes from host movements relies on the key assumption that environmental characteristics strongly influence movement and space use. While there is a growing number of methods that quantify the effect of environmental characteristics on movement and habitat use (reviewed in Hooten et al., 2017), there has been much less work exploring how (and if) environment affects and predicts correlation and synchrony among hosts. Therefore, an essential next-step for building epidemiological landscapes will be predictive models of spatio-temporal correlation among individuals (e.g. Brandell et al., 2021). Our analyses emphasize the importance of this next step by showing that correlations at a local (cell) scale can create a vastly more heterogeneous transmission landscape. Moreover, correlations can lead to localized transmission hotspots that are not necessarily predictable from joint space use (Yang et al., 2023a), and may contribute to the growing empirical recognition that fine-scale, localized transmission hotspots are present in many empirical host-pathogen systems (Albery et al., 2021). Whether these localized transmission hotspots are predictable *a priori* remains to be seen, but ignoring correlated movements can make the task orders of magnitude more difficult for some pathogens.

## Acknowledgments

Thanks to Dr. A. Houston, B. Trout, M. Turner and TWRA for capture and logistical support during fieldwork. Funding for this study was provided by the United States Department of Agriculture (USDA) National Institute of Food and Agriculture (McIntire Stennis project TEN00MS-113; Hatch Project 7001607) and the USDA American Rescue Plan (APP-21535). A portion of the computation for this work was performed on the University of Tennessee Infrastructure for Scientific Applications and Advanced Computing (ISAAC) computational resources.

