## Appendix for "Correlated host movements can reshape spatio-temporal disease dynamics: modeling the contributions of space use to transmission risk using movement data"

### Appendix 1: PMoveSTIR in continuous space

In the main text, we derive PMoveSTIR assuming that hosts are moving and contacting each other within some area $A_x$, which we can conceptually think about as a grid cell on a gridded landscape. While this conceptually simplifies the problem, it is more general to consider the case of continuous space where we define contact as potentially happening when present or past host $j$ is (was) within some distance $r$ of host $i$ at its present location (Wilber et al., 2022). The output we want from this alternative version of PMoveSTIR is the function $\hat{h^*}(x,t)$, which is the force of infection *per unit area* at point $x$ on the landscape (e.g., $\hat{h^*}(x,t)$ might have units day$^{-1}$m$^{-2}$). Integrating this function over different areas will yield estimates of force of infection felt by host $i$ from host $j$ for any area of interest on the landscape.

Let's start with the situation where a host $i$ is occupying some circular area $A_{x,\rho}$ where $x$ is the center of the area and $\rho$ is the radius of the area. A contact can occur when host $j$ (past or present) is in the area $A_{x,\rho+r}$ where $r$ is our epidemilogically relevant contact distance and $r >> \rho$. The force of infection felt by host $i$ from host $j$ as time $t$ in area $A_{x,\rho}$ is given by

$$h_{i \leftarrow j}(t, A_{x,\rho}) = \int_{-\infty}^{t} \beta' \lambda \delta'_{x_i(t)}(A_{x,\rho}) \delta'_{x_j(u)}(A_{x,\rho+r}) e^{-\nu(t-u)} du. \tag{1}$$

where, consistent with the main text, $\delta'_{x_i(t)}(A_{x,\rho})$ is a Bernoulli random variable that determines whether or not host $i$ is located in area $A_{x,\rho}$ at time $t$ and $\delta'_{x_j(u)}(A_{x,\rho+r})$ is a Bernoulli random variable that determines whether or not host $j$ is in area $A_{x,\rho+r}$ at time $u$. The variables $x_i(t)$ and $x_j(u)$ indicate the locations of host $i$ and $j$ at time $t$ and $u$, respectively. The parameter $\beta' = \frac{\tilde{\beta}}{A_{x,\rho+r}}$ and indicates our assumption that encounters are equally likely within an area $A_{x,\rho+r}$. The parameter $\lambda$ is the pathogen shedding rate of host $j$ and $\nu$ is the pathogen decay rate once deposited in the environment. As in the main text, we are computing maximum transmission risk and assuming that host $j$ is always infected at any time $u$.

As we did in the main text, we can envision simulating many different movement trajectories for host $i$ and $j$ and take the expectation of $h_{i \leftarrow j}(t, A_{x,\rho})$. We obtain

$$h^*_{i \leftarrow j}(t, A_{x,\rho}) = \int_{-\infty}^{t} \beta' \lambda E[\delta'_{x_i(t)}(A_{x,\rho}) \delta'_{x_j(u)}(A_{x,\rho+r})] e^{-\nu(t-u)} du. \tag{2}$$

We can rewrite equation 2 as

$$h^*_{i \leftarrow j}(t, A_{x,\rho}) = \frac{\tilde{\beta}}{A_{x,\rho+r}} \lambda \int_{-\infty}^{t} [p_i(A_{x,\rho}, t) p_j(A_{x,\rho+r}, u) + Cov(\delta'_{x_i(t)}(A_{x,\rho}), \delta'_{x_j(u)}(A_{x,\rho+r}))] e^{-\nu(t-u)} du, \tag{3}$$

where the force of infection felt by host $i$ from host $j$ is the related to the utilization distributions of the two hosts and their covariance within an area (in actuality, two nested areas).

For simplicity, let's consider hosts moving independently. In this case, we can write

$$h^*_{i\leftarrow j}(t, A_{x,\rho}) = \beta' \lambda \int_{-\infty}^{t} [p_i(A_{x,\rho}, t) p_j(A_{x,\rho+r}, u)] e^{-\nu(t-u)} du \tag{4}$$

where $p_i(A_{x,\rho}, t)$ is the probability of host $i$ being in area $A_{x,\rho}$ at time $t$ and $p_j(A_{x,\rho+r}, u)$ is the probability of host $j$ being in area $A_{x,\rho+r}$ at time $u$.

Now we want to calculate $h^*_{i\leftarrow j}(t, A_{x,\rho})$ in the limit as $\rho \to 0$. For equation 4, we can divide both sides by $A_{x,\rho}$ and take the limit as $\rho \to 0$ (such that $A_{x,\rho} \to 0$). Doing this we obtain

$$\hat{h}^*_{i\leftarrow j}(t, x) = \beta' \lambda \int_{-\infty}^{t} [f_i(x, t) p_j(A_{x,r}, u)] e^{-\nu(t-u)} du = \beta' \lambda \int_{-\infty}^{t} [f_i(x, t) \int_{s \in A_{x,r}} f_j(s, u) ds] e^{-\nu(t-u)} du \tag{5}$$

where $f_i(x, t)$ and $f_j(x, t)$ are the probability density functions of space use for host $i$ and host $j$, respectively. Note that the units on $f_i(x, t)$ or $f_j(x, t)$ are per area, such that the force of infection $\hat{h}^*_{i\leftarrow j}(t, x)$ has units per time per area as opposed to $h^*_{i\leftarrow j}(t, A_{x,\rho})$ which has units per time. Conceptually, for $h^*_{i\leftarrow j}(t, A_{x,\rho})$ we have already integrated over area so we cancel out the per area units.

Assuming a stationary process, we can write equation 5 as

$$\hat{h}^*_{i\leftarrow j}(t, x) = \frac{\beta' \lambda}{\nu} [f_i(x) \int_{s \in A_{x,r}} f_j(s) ds] \tag{6}$$

Furthermore, if we assume that the space use of host $j$ is relatively uniform within the contact area $A_{x,r}$ we can simplify to

$$\hat{h}^*_{i\leftarrow j}(t, x) = \frac{\beta' \lambda}{\nu} [f_i(x) f_j(x) \pi r^2] \tag{7}$$

Remembering that $\beta' = \tilde{\beta}/A_{x,r} = \tilde{\beta}/\pi r^2$, we get

$$\hat{h}^*_{i\leftarrow j}(t, x) = \frac{\tilde{\beta} \lambda}{\nu} [f_i(x) f_j(x)] \tag{8}$$

Integrating $\hat{h}^*_{i\leftarrow j}(t, x)$ over some area of interest centered at $x$ would yield $h^*_{i\leftarrow j}(t, A_{x,d}) = \frac{\tilde{\beta}\lambda}{\nu} \int_{A_{x,d}} [f_i(s) f_j(s) ds]$. This is reminiscent of the equation 17 in Martinez-Garcia et al. (2020) where a limiting case of the mean encounter rate of two individuals moving according to an Ornstein-Uhlenbeck movement process is proportional to the inner product of their utilization distributions.

Including correlation in movement into this formulation of PMoveSTIR is more theoretically and empirically challenging, and we leave this task for a later paper.

### Appendix 2: Deriving PMoveSTIR given an assumption of statistical stationarity

To derive equation 5 in the main text that assumes stationarity in utilization distributions, we start with equation 4 in the main text

$$h^*_{i \leftarrow j}(t, x) = \frac{\tilde{\beta}}{A_x} \lambda \int_{-\infty}^{t} [p_i(x, t) p_j(x, u) + Cov(\delta'_{x_i(t)}(x), \delta'_{x_j(u)}(x))] S(t - u) du. \tag{9}$$

Here, $p_i(x, t)$ and $p_j(x, u)$ represent the probabilities of host $i$ and $j$ using location $x$ at time $t$ and $u$, respectively. The parameters $\tilde{\beta}$ and $\lambda$ are the acquisition and deposition rates respectively, and $A_x$ is the area of location $x$. $Cov(\delta'_{x_i(t)}(x), \delta'_{x_j(u)}(x))$ gives the covariance in how host $i$ and $j$ at time $t$ and $u$ are using area $x$. Finally, $S(t - u)$ gives the survival probability of a pathogen at time $t$ that was deposited at time $u$.

If we now assume a stationary utilization distribution, the time indexes on $p_i(x, t)$ and $p_j(x, u)$ are irrelevant – the logic here is that, by definition, the mean of a stationary distribution is independent of time so $p_i(x, t) = p_i(x)$ and $p_j(x, u) = p_j(x)$. Therefore, we can write

$$h^*_{i \leftarrow j}(t, x) = \frac{\tilde{\beta}}{A_x} \lambda \left[ p_i(x) p_j(x) \int_0^{\infty} S(\tau) d\tau + \int_{-\infty}^{t} Cov(\delta'_{x_i(t)}(x), \delta'_{x_j(u)}(x)) S(t - u) du \right].$$

In addition, given stationarity, $h^*_{i \leftarrow j}(t, x)$ does not depend on time, such that

$$h^*_{i \leftarrow j}(x) = \frac{\tilde{\beta}}{A_x} \lambda \left[ p_i(x) p_j(x) \int_0^{\infty} S(\tau) d\tau + \int_{-\infty}^{t} Cov(\delta'_{x_i(t)}(x), \delta'_{x_j(u)}(x)) S(t - u) du \right].$$

Moreover, given stationarity, we know that $Cov(\delta'_{x_i(t)}(x), \delta'_{x_j(u)}(x))$ just depends on the time lag $s = t - u$ and not specific time stamps. Thus, we can write

$$h^*_{i \leftarrow j}(x) = \frac{\tilde{\beta}}{A_x} \lambda \left[ p_i(x) p_j(x) \int_0^{\infty} S(s) d\tau + \int_0^{\infty} Cov(\delta_{i \in x}, \delta_{j \in x} | \tau) S(s) ds \right]. \tag{10}$$

By replacing $S(s)$ with the exponential survival function $e^{-\nu s}$ we obtain equation 5 in the main text.

In the lower left-hand corner of Fig. 1 in the main text, we have the case where space use is uniform and time is stationary. We again assume that pathogen decay is exponentially distributed with a rate of decay

$\nu$ such that $S(s) = \exp(-\nu s)$. Given these assumptions, we can write equation 10 as

$$h^*_{i \leftarrow j}(A_x) = \beta' \lambda \left[ \frac{A_x}{A_{tot}} \frac{A_x}{A_{tot}} \frac{1}{\nu} + \int_0^\infty Cov(\delta_{i \in A_x}, \delta_{j \in A_x} | s) e^{-\nu s} ds \right] \quad (11)$$

where the covariance in contact is constant across all areas $A_x$ on the landscape (such that $\delta_{i \in A_x}$ indicates the use of some arbitrary area $A_x$). If hosts are moving independently (i.e., covariance is 0) we obtain $\frac{\tilde{\beta}}{A_x} \frac{A_x}{A_{tot}} \frac{A_x}{A_{tot}} \frac{\lambda}{\nu}$. Given a gridded landscape with non-overlapping grids and $x$ is a single grid cell, summing over all $n$ areas $A_x$ that comprise the landscape yields $\bar{h}_{i \leftarrow j} = \frac{\tilde{\beta}}{A_{\text{tot}}} \frac{\lambda}{\nu}$, which is the standard mass action assumption of transmission (McCallum, 2001).

### Appendix 3: Beyond the corners of PMoveSTIR

PMoveSTIR also accounts for other useful cases regarding how heterogeneity in space and time relate to the expected FOI. For example, home range overlap is an intermediate case of of PMoveSTIR (Fig. 1 in the main text), where time is stationary, space use is uniform within a home range, and the area of contact is the home range overlap. Transmission networks commonly assume that this overlap is proportional to the edge weight between host $i$ and host $j$ (e.g. Springer et al., 2017). Given an area of overlap $A_{hro}$ between two individual home ranges with areas $A_{tot,i}$ and $A_{tot,j}$, we can write

$$h^*_{i \leftarrow j}(A_{hro}) = \frac{\tilde{\beta}}{A_{hro}} \lambda \left[ \frac{A_{hro}}{A_{tot,i}} \frac{A_{hro}}{A_{tot,j}} \frac{1}{\nu} + \int_0^\infty Cov(\delta_{i \in A_{hro}}, \delta_{j \in A_{hro}} | s) e^{-\nu s} ds \right] \quad (12)$$

which gives us an explicit equation for how home range overlap determines FOI and how correlation in use of the overlap area affects FOI.

In many cases, animals shift their home range seasonally (Viana et al., 2018; Richard et al., 2014) such that assuming a constant UD is biologically unrealistic. This can be accounted for in PMoveSTIR as a special case of temporal heterogeneity (Fig. 1 in the main text). Specifically, consider equation 3 in the main text. The expected FOI over the time interval 0 to $t$ in location $x$ is $\bar{h}^*_{i \leftarrow j}(x) = \frac{\int_0^t h^*_{i \leftarrow j}(\tau, x) d\tau}{t}$ (Wilber et al., 2022). One could approximate this as $\sum_{k=0}^{n_t} h^*_{i \leftarrow j}(t_k, x) \frac{\Delta \tau}{t}$ where $n_t$ is the number of bins that comprise the interval 0 to $t$, $t_k$ is the midpoint of the $k$th time interval, and $\Delta \tau$ is the width of a time interval. The equation is a weighted sum where each component is a constant FOI within a time interval multiplied by the length of the time interval relative to the total interval length $t$.

The summation perspective helps us frame PMoveSTIR in a seasonal context. For example, consider that hosts use two primary movement patterns that repeat on a period of $T$ (e.g., over the course of a year).

The first movement pattern lasts for $\tau_1$ time units and the second lasts for $\tau_2$ time units where $\tau_1 + \tau_2 = T$. Assuming stationarity within each time interval, the average FOI felt by host $i$ from host $j$ in location $x$ over period $T$ is

$$\bar{h}^*_{i\leftarrow j}(x) = \frac{\tau_1}{T} h^*_{\tau_1, i\leftarrow j}(x) + \frac{\tau_2}{T} h^*_{\tau_2, i\leftarrow j}(x) \tag{13}$$

This could be generalized to any number of intervals within the period $T$, depending on the biology of the system. Importantly, when considering the epidemiological implications of these seasonally shifting movement patterns, one should consider each seasonal FOI component separately to build dynamic contact networks with time-varying edges (Wilber et al., 2022).

### Appendix 4: Analytical example of how correlated movement and indirect transmission can alter FOI

The effects of correlated movement on FOI relative to habitat overlap become analytically more difficult to generalize when we consider pathogens with indirect transmission. This is because the correlation function $Cor(\delta_{i\in A_x}, \delta_{j\in A_x}|s)$ can be highly non-trivial even in the simple case when two hosts are moving together (see Fig. 1 for some examples). While determining the analytical form of $Cor(\delta_{i\in A_x}, \delta_{j\in A_x}|s)$ for common movement models is beyond the scope of this study, we can use a relatively simple movement scenario to get an analytical sense of how indirect transmission and correlated movements can interact to affect transmission risk.

The scenario is two hosts moving back and forth between two patches, and spending $\eta$ time units in each patch. As with the example of direct transmission in the main text, we assume that the hosts are moving together. If we plot $Cor(\delta_{i\in A_x}, \delta_{j\in A_x}|s)$ of this movement scenario, we get the saw-tooth pattern observed in Fig. 1a. Correlation in movement is 1 at lag 0 (because hosts are together constantly) and decreases to approximately -1 by time $\eta$ before increasing close to 1 by time $2\eta$, etc. We assume that the pathogen cannot survive longer than $\eta$ in the environment (i.e., $\pi \leq 1$), then we can ignore all correlations after lag $s = \eta$ and we can approximate $Cor(\delta_{i\in A_x}, \delta_{j\in A_x}|s) \approx (-2/\eta)s + 1$. With this approximation, we can write our FOI equation as

$$h^*_{i\leftarrow j}(x) = \beta' \lambda \pi \eta \left[ \underbrace{\frac{A_x}{A_{tot}} \frac{A_x}{A_{tot}}}_{\substack{\text{Contribution due}\\ \text{to habitat overlap}}} + \underbrace{\frac{A_x}{A_{tot}} (1 - \frac{A_x}{A_{tot}})(1 - \pi)}_{\substack{\text{Contribution due}\\ \text{to correlated movement}}} \right]. \tag{14}$$

As $\pi$ gets small, we recover equation 7 in the main text. Equation 14 shows that the degree of contribution of correlated movement to FOI depends on the relative persistence time of the pathogen compared to the residence time of the host. When the pathogen persists from $\eta$ time units (i.e. $\pi = 1$) the contribution of correlation to FOI is zero. This is because negative and positive correlations in $Cor(\delta_{i\in A_x}, \delta_{j\in A_x}|s)$ cancel out as we integrate from zero to $\eta$. When $\pi < 1$, the contribution of correlated movements to FOI will be positive, and the relative contribution of correlated movements compared to habitat overlap will approach unity (i.e., an equal contribution) as $\pi$ approaches 0. This example illustrates three important points: the contributions of correlated movement to indirect transmission risk will depend strongly on 1) the movement dynamics as reflected in $Cor(\delta_{i\in A_x}, \delta_{j\in A_x}|s)$ and 2) the rate of pathogen decay relative to host movement rates. Third, the contribution due to correlated movements could potentially increase, decrease, or have no effect on local FOI depending on the lag considered.

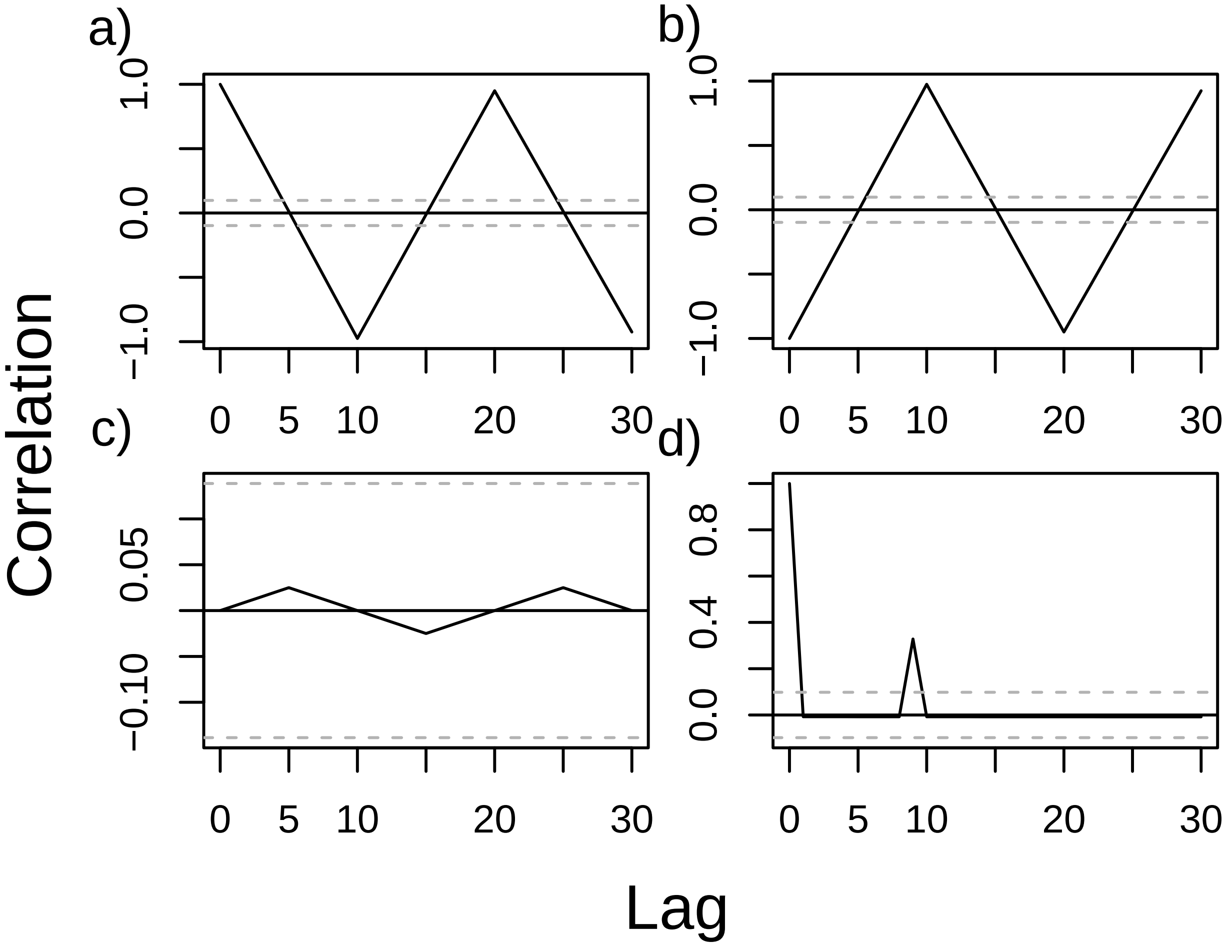


Figure 1: Variation in cross-correlation values for different combinations of position histories. The extreme variety of cross-correlation patterns between two moving hosts makes the effect of correlation on indirect transmission more difficult to generalize than direct transmission.
